# Sample-specific CpG loci are important for accurate long-read methylation analysis

**DOI:** 10.1101/2025.04.30.651558

**Authors:** Søren B. Hansen, Kasper D. Hansen, Morten T. Limborg

## Abstract

Methylated CpG loci mutate at high rates, but it remains challenging to identify sample-specific non-reference CpG loci for methylation analyses. Here, we present a likelihood-based method that identifies sample-specific CpG loci at high precision in long-read sequencing data from Nanopore and PacBio platforms. Inclusion of non-reference CpG loci increased the number of loci and reduced reference bias, and is important for accurate estimation of global methylation levels across samples.

## Main

In vertebrate genomes, cytosines in CpG context are frequently modified by a methyl group on the 5th carbon position (5mC)^1^ and methylation profiles are increasingly studied for their evolutionary importance in adaptive phenotypes^2^. Methylated cytosine has a high mutation rate, and often a discrepancy between the position of CpG loci in a sample and the reference genome challenges analysis and interpretation^3^. In data from conversion based methods, such as bisulfite sequencing, it is necessary to use personal reference genomes to avoid reference bias^4^. This approach is used in controlled laboratory lines with high quality reference genomes^5^, but for ecologically relevant wild samples a more feasible solution is to remove CpG loci if they overlap known genetic variation^6^. Removing polymorphic CpC loci in the sample set effectively reduces reference bias, however, it requires genotyping the sample set and further removes sample-specific CpG loci, which may be especially important to study in an evolutionary context. Single-molecule long-read sequencing of DNA on sequencing platforms from Pacific Biosciences (PacBio) and Oxford Nanopore Technologies (Nanopore) enables detection of methylation directly from the raw sequencing signal^7,8^. The simultaneous sequencing of nucleotides and methylation enables identification and analysis of non-reference CpG loci, without prior genotyping or personal reference genomes. Here, we present a precise likelihood-based framework to identify sample-specific CpG loci in long-read data from both PacBio and Nanopore platforms with benefits for downstream analysis. The framework has been integrated in the R-package BSseq^9^.

We validated our approach using publicly available, long-read data from the GIAB sample, HG002, which is accompanied by high-quality variant calls, when mapped to the GRCh38 genome assembly^10,11^. On chromosome 1 used for benchmarking, the mean coverage was 37X in the Nanopore sample^12^ and 20X in the PacBio sample^13^ (Fig. 1a). The number of reference loci with at least one CpG site mapped was 3,041,114 in the Nanopore sample, and 2,195,403 in the PacBio sample, including only the high confidence regions^14^. This exceeded the number of reference CpG loci (2,132,501), but included many loci with a high likelihood of being non-CpG where the mapped CpG site was caused by sequencing or mapping errors (Fig. 1b). The two sequencing technologies had similar error rates for miscalling true CpG sites as non-CpG sites, despite PacBio having an overall lower error rate across all dinucleotides. Loci with a scaled likelihood higher than 99% of being homozygous CpG (.99 homozygous filtered), or heterozygous CpG (.99 heterozygous filtered) showed high concordance with the genotype of the HG002 sample. In both the Nanopore and PacBio sample, the precision of the .99 filtered loci exceeded 99%, when compared with a truth set of homozygous and heterozygous reference CpG loci. The precision was equally high (>98%), when using a truth set of homozygous and heterozygous non-reference CpG loci (Extended data: Table 1 and Table 2). The main challenge was to distinguish homozygous and heterozygous CpG loci, wherefore the precision, and the number of loci increased, when we filtered for a combined scaled likelihood higher than 99% of being homozygous or heterozygous (.99 all filtered) (Fig. 1c, Extended data: Table 1 and Table 2). The .99 filtered data was largely overlapping between the Nanopore and Pacbio sample and the loci obtained from joint .99 filtering in both samples had precision above 99.9% in all comparisons (Fig. 1d, Extended data: Table 3). These loci, called at high precision included 15,040 joint .99 homozygous filtered loci, 21,034 joint .99 heterozygous filtered loci, and 39,017 joint .99 all filtered CpG loci not overlapping reference CpGs. This corresponded to 0.8% of all homozygous CpG loci, 46.0% of all heterozygous CpG loci, and 1.8% of the all filtered loci in the sample, which would have been lost using reference-guided filtering (Extend data: Table 4).

**Fig 1.**
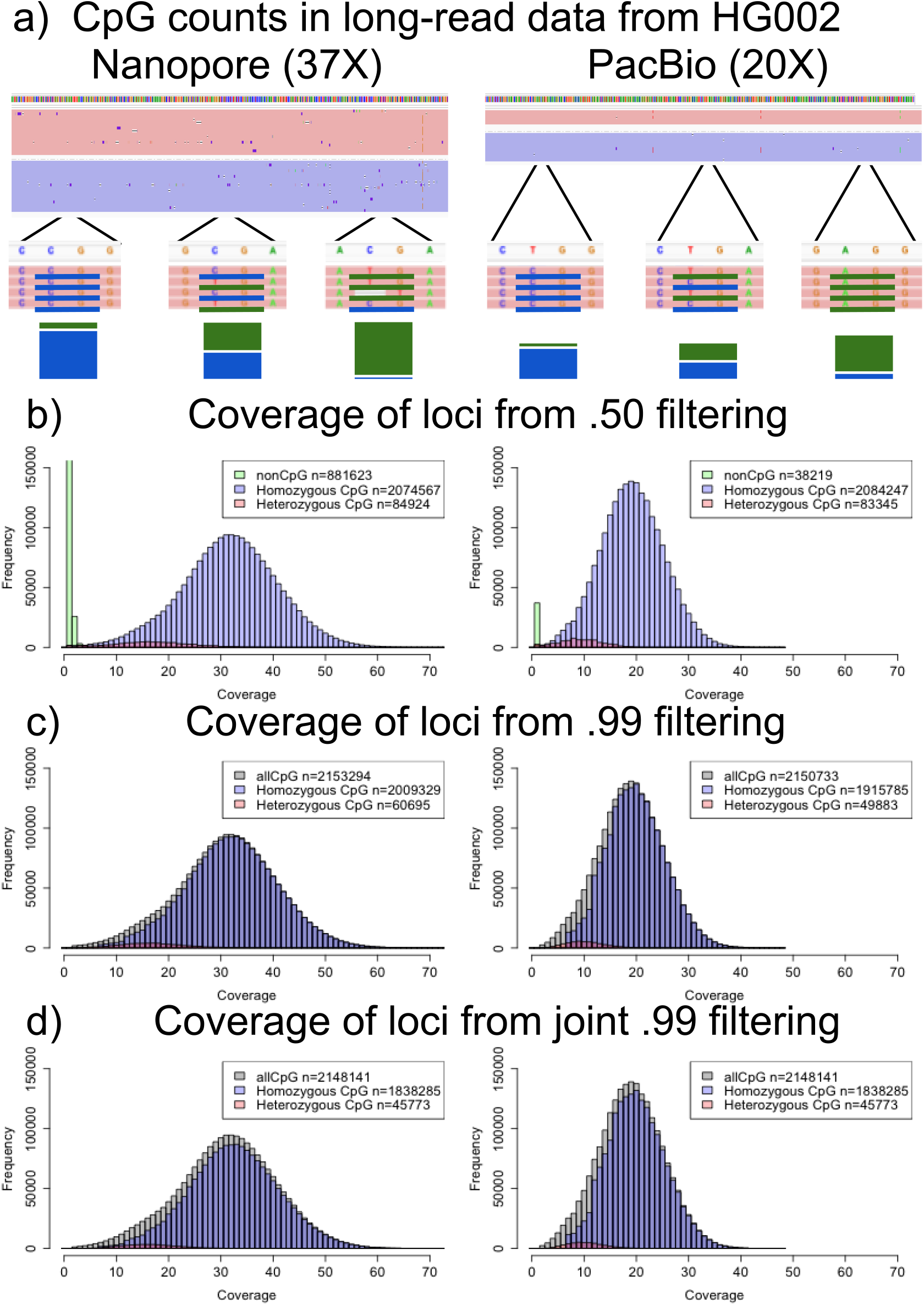
Likelihood method identifies sample-specific CpG loci in long-reads data. **a)** The count of CpG sites and non-CpG sites for each reference loci is counted in HG002 long-read data from Oxford Nanopore (left) or PacBio (right) platforms mapped to GRCh38. **b-d)** Coverage of loci colored by the most likely CpG status given the count data: Homozygous CpG (blue), heterozygous CpG (red), non-CpG (green), all CpG (grey) using a.50 threshold **(b)**, a .99 threshold **(c)**, and a joint .99 threshold in both samples **(d)**.

To demonstrate the importance of including non-reference CpG loci in a challenging, but ecologically and evolutionarily relevant sample set, we analysed nine fish from the species, Atlantic silversides (*Menidia menidia*). Atlantic silversides have one of the highest levels of heterozygosity reported in any vertebrate, including structural variation such as a 9.4 Mb inversion on the 17.2 Mb long chromosome 24^15^. The inversion suppresses recombination, and there is high genomic divergence between the homologous DNA sequences within the inversion in both wild and experimental individuals^16–18^. We generated Nanopore data from nine fish with different chromosome 24 inversion configurations; having either two reference-like (RR) (n=3), one reference-like and one non-reference-like (RN) (n=3) or two non-reference-like (NN) (n=3) versions of the inversion orientation (Fig. 2a-b). All nine samples were mapped to the same reference genome, and the mean coverage of chromosome 24 was 31X per sample. Using joint .99 all filtering for each inversion configuration and only the loci which overlapped reference CpG loci (reference-guided filtering), the highest number of loci was retained in the RR samples (304,231), with 1.5% fewer loci in the RN samples, and 7.4% fewer loci in the NN samples (Extend data: Table 5). The density of the reference-guided filtered loci (loci per 100kb) was lower within the inversion in the NN samples, suggesting a reference bias (Fig. 2c). When non-reference CpG sites were included, the highest number of CpG loci were identified in the RN samples (327,821) (Extend data: Table 5). Between the NN samples and the RR samples, the joint .99 filtering removed the systematic difference in CpG loci density within the inversion, and reduced the difference in the number of CpG loci from 7.4% to 1.2% (Fig. 2d, Extend data: Table 5). Combined, .99 all filtering retained 4.3% more CpG loci in the RR samples, 8.6% more loci in the RN samples, and 10.1% more loci in the NN samples by including both homozygous and heterozygous non-reference loci (Fig. 2e-f, Extend data: Table 5).

**Fig. 2:**
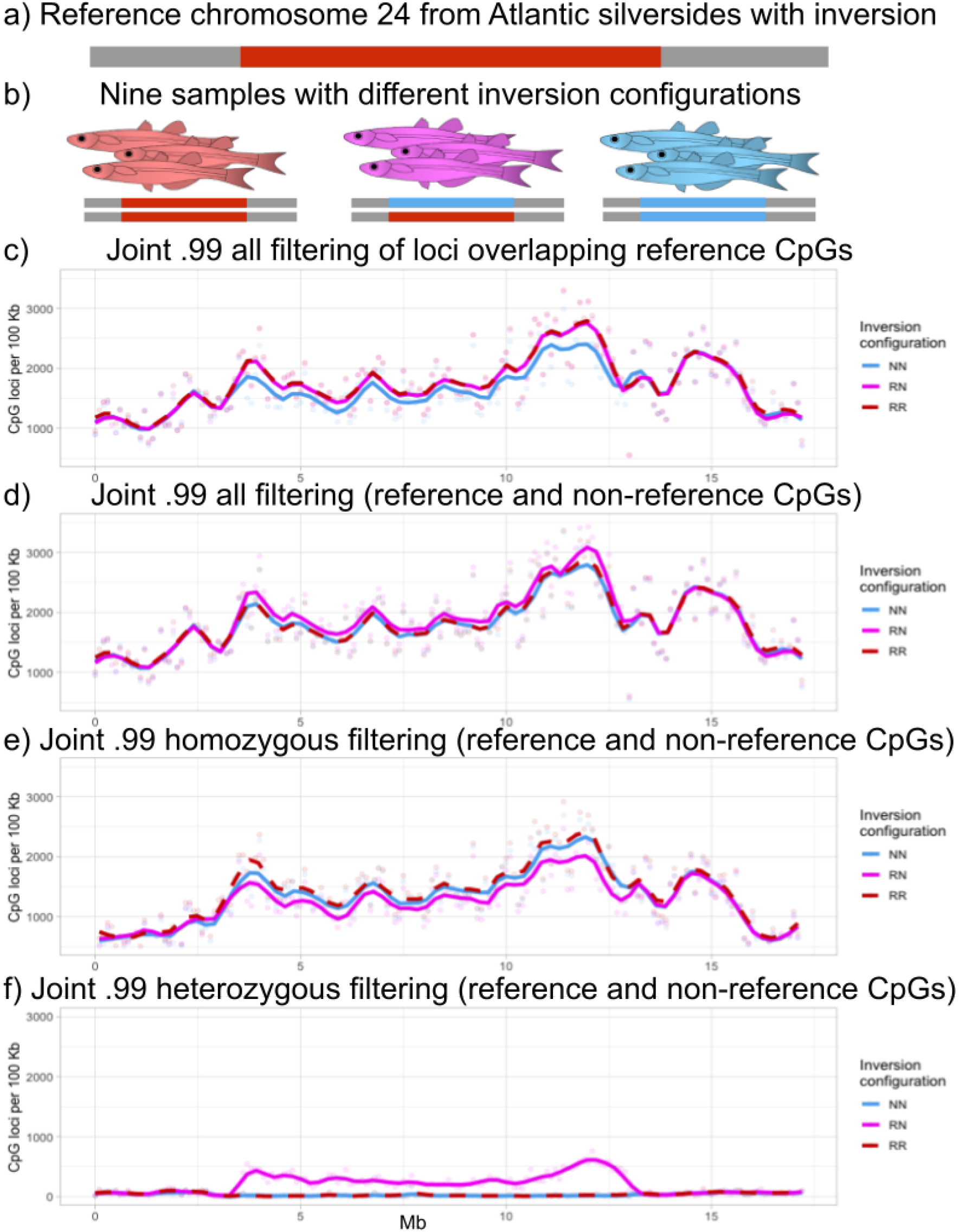
Likelihood filtering removes reference bias in challenging samples. **a)** Reference chromosome 24 with reference-like (R) inversion orientation. **b)** Three samples homozygous for the reference-like orientation (RR), three samples heterozygous for reference-like and non-reference-like inversion orientation, (RN) and three samples homozygous for the non-reference-like orientation (NN). **c-f)** Number of: joint .99 all filtered reference CpG loci **(c)**, joint .99 all filtered loci **(d)**, joint .99 homozygous filtered loci **(e)**, and joint .99 heterozygous filtered loci **(f)** per 100 KB in the three RR samples in red, the three NN samples in blue and all three RN samples in purple.

In addition to increasing the number of loci for downstream analysis, likelihood filtering enables accurate estimation of global methylation levels. In the three different chromosome configurations in the Atlantic silverside samples, the mean chromosome-wide methylation level of the .99 all filtered non-reference CpG loci were respectively 5.8 percentage points (*%pt*.) higher in the RR samples, 3.7 *%pt*. higher in the RN samples, and 3.4 *%pt*. higher in the NN samples, compared to the mean methylation level of the reference CpG loci (Extend data: Table 6). In the HG002 sample, the mean methylation of the joint .99 all filtered non-reference CpG loci was 2.3 *%pt*. higher in the Nanopore sample, and 0.9 *%pt*. higher in the PacBio sample, compared to the reference CpG loci (Extend data: Table 7). The higher methylation level in non-reference CpG loci was confounded by a higher methylation level in heterozygous than homozygous CpG loci.

Our method efficiently identifies sample-specific CpG loci whereas reference-guided filtering of a diploid sample using a haploid reference genome removes half of the heterozygous CpG loci and a number of homozygous CpG loci, depending on the evolutionary distance between the sample and the reference genome. While genotyping samples prior to a methylation analysis and/or using diploid reference genomes or pan genomes, may also recover sample-specific CpG loci, we offer a convenient alternative using only standard methylation output. It remains a challenge to compare sample-specific CpG loci in downstream analyses. Contrary, focusing exclusively on the CpG loci overlapping between all samples and the reference genome may be a barrier for understanding adaptive methylation over evolutionary times.

## Methods

### Analysis of Nanopore and PacBio data from HG002

We downloaded publicly available Nanopore^12^ and PacBio^13^ data from the HG002 sample of the GIAB project. The pod5 files in the pass directory from the Nanopore sample was modification and base called using the model: dna_r10.4.1_e8.2_400bps_sup@v4.3.0_5mCG_5hmCG@v1 from dorado/0.6.0^19^, while the downloaded PacBio reads were modification and base called using an on-instrument HiFi base and 5mCpG model. The bam files with modification and base called reads were mapped against the human genome assembly GRCh38^11^ using minimap2/2.26^20^ and samtools/1.20^21^ and the coverage was estimated using samtools depth. The Nanopore modification calls were filtered using default settings of Modkit pileup to produce a bedMethyl file. For piling up the PacBio sample we set the flag --no-filtering to avoid warnings because of different quality encoding when producing the bedMethyl file in Modkit/0.4.2^22^. The specific version of each software is not important to get the desired output likelihood filtering, but it is a strict necessity that the modification and base calling is performed using a CpG context model, and the pileup is not using reference-guided filtering (i.e. no --cpg or --traditional preset flag) (Extended data: Fig. 1).

The bedMethyl files from both samples were imported into BSseq v.1.45.1 as BSseq objects using read.bedMethyl() and we filtered for loci in high confidence regions with no inconsistent variant calls^14^. The CpG specific error rate was estimated to be 3.10% for the Nanopore sample and 3.24% in the PacBio sample, although fewer non-CpG sites seemed misclassified as CpG sites in the PacBio sample (Extended data: Fig. 2). We used .50 filtering of homozygous loci, heterozygous loci and non-CpG loci in the Nanopore and PacBio sample using getCpG() function setting type to “homozygous”, “heterozygous”, or “nonCpG”, and the threshold to 0.50. This filtering resulted in respectively 2,074,567 homozygous, 84,924 heterozygous and 881,623 non-CpG loci from the 3,041,114 loci with a coverage of at least one in the Nanopore sample and 2,084,247 homozygous, 83,345 heterozygous and 38,219 non-CpG loci from the 2,195,403 loci with a coverage of at least one in the PacBio sample. The coverage of these loci colored by type were plotted in Fig. 1b.

Secondly, we used a more conservative .99 filtering to obtain homozygous loci, heterozygous loci, and homozygous or heterozygous loci, using getCpG() function setting type to “homozygous”, “heterozygous” or “allCpG”, and the threshold to 0.99. This filtering resulted in 2,009,329 homozygous, 60,695 heterozygous and 2,153,294 all filtered loci in the Nanopore sample, and 1,915,785 homozygous, 49,883 heterozygous and 2,150,733 all filtered loci in the PacBio sample. The coverage of these loci colored by type were plotted in Fig. 1c.

Lastly, we constructed a matrix of the most likely CpG status and the corresponding scaled likelihood from the multi-sample BSseq object containing both the Nanopore and PacBio sample using getCpGMatrix() and getMaxLikelihoodMatrix(). Using these matrices, we jointly filtered the BSseq object for homozygous loci, heterozygous loci and allCpG loci with a scaled likelihood higher than 0.99 in both samples. This filtering resulted in 1,838,285 homozygous, 45,773 heterozygous and 2,148,141 all filtered loci. The coverage of these loci colored by type were plotted in Fig. 1d.

### Validation using truth set of homozygous, heterozygous, and non-CpG loci

The HG002 sample is accompanied by a “truth” set of highly consistent variants called across several sequencing platforms at high quality. We downloaded a VCF file of these variants along with the file specifying high-confidence regions of the GRCh38 reference genome with no ambiguous calls^10,14^. We identified all CpG loci and CH loci on both strands of the reference chromosome 1 using Modkit motif, and converted the positions to match the BSseq object. We generated a truth set of homozygous reference CpG loci, by identifying reference CpG loci not overlapping variants in the VCF file. A truth set of heterozygous reference CpG loci was made by identifying heterozygous C to D and G to H variants overlapping reference CpG loci. Correspondingly, a truth set of reference non-CpG loci was generated from non-CpG loci in the reference chromosome not overlapping variants in the VCF file. We generated a truth set of homozygous non-reference CpG loci and heterozygous non-reference CpG loci, from homozygous, and heterozygous H to G or D to G variants overlapping CH loci adjusting for variant and strand. The precision of the .99 filtered homozygous loci, the .99 filtered heterozygous loci and the .99 all filtered loci was estimated by evaluating the number of correctly or incorrectly classified loci in the truth sets (Extended data: Table 1, Table 2 and Table 3). We used the reference CpG loci to distinguish between reference and non-reference CpG loci in the BSseq objects, and evaluated the overlap with the truth sets, which only included CpG loci introduced by SNPs (Extended data: Table 4).

### Analysis of data from Atlantic silversides with complex genetic structure

For an ecological and evolutionary relevant sample, we tested our method on the small annual pelagic fish, the Atlantic Silversides (*Menidia menidia*). Briefly, the data was generated from DNA extracted from the gill tissue of nine samples, using the Nanopore R10.4.1 long-read sequencing service from Novogene (UK). The modification and base calling was performed using the same model as for the HG002 Nanopore sample, and the reads were mapped to the Atlantic Silverside reference genome^15^. The mean estimated coverage of chromosome 24 was 31X (standard deviation = 6X) estimated using samtools depth. Similar to the HG002 Nanopore sample, the bedMethyl files were produced using Modkit pileup with default filtering of low quality modification calls. The files from the nine samples were read as a multisample BSseq object using read.bedMethyl(). We filtered for loci on chromosome 24 and estimated the CpG specific error rate to be in the range of 3.4% to 3.8% per sample. The CpGMatrix and the MaxLikelihoodMatrix was constructed using getCpGMatrix() and getMaxLikelihoodMatrix() in a normal version and an allCpG version by setting AllCpG=T. To compare the .99 filtered results to reference-guided filtering, we identified CpG sites in the reference genome using Modkit motif.

We performed joint filtering of the samples based on *a priori* information on their chromosome 24 inversion configuration. The BSseq object and the corresponding CpGMatrices and MaxLikelihoodMatrices was subset to contain the three samples with either RR, RN or NN chromosome 24 configuration of inversion orientations (Fig. 2a). Firstly, each of the three BSseq objects (multi-sample objects containing either three RR samples, three RN or three NN samples) was joint .99 all filtered using the allCpG version of the CpGMatrix and MaxLikelihoodMatrix. Secondly each of the three BSseq objects were joint .99 homozygous filtered, or joint .99 heterozygous filtered using the normal version of the CpGMatrix and MaxLikelihoodMatrix. We counted the overlap and difference between the .99 filtered loci and the reference CpG loci (Extended Data: Table 5), and plotted the density of the .99 filtered loci along the chromosome using ggplot2 (Fig. 2c-f). The described filtering is based on prior knowledge of the relevant genetic structure in the sample set. In sample sets where the genetic structure is expected to be important but unknown, the CpGMatrix can serve as a convenient input for analyses such as PCA to resolve genetic structure (data not shown).

### Estimating mean global methylation levels

The filtered BSseq objects included the summarized modification information from the long-read data. We calculated the mean of the beta values from getMeth() of the BSseq objects setting type to “raw”. For each of the nine silverside samples, we computed the mean of reference CpG and non-reference CpG loci from the joint .99 all filtered loci, the joint .99 homozygous filtered loci and the joint .99 heterozygous filtered loci (Extended data: Table 6). We made a similar comparison of the mean methylation levels of reference CpG and non-reference CpG loci in both the Nanopore and PacBio data from the joint .99 filtered HG002 sample (Extended data: Table 7). Although the loci of the two HG002 samples overlap, the methylation estimates are based on different technologies, different filtering, and different handling of 5hmC calls, wherefore they are not comparable.

## Data and code availability

The Nanopore sequencing data in pod5 format was downloaded from the run id: 20230424_1302_3H_PAO89685_2264ba8c from https://epi2me.nanoporetech.com/giab-2023.05/ (last checked 25/4 -2025) and the PacBio data was downloaded in bam format from https://downloads.pacbcloud.com/public/2024Q4/Vega/HG002/ (last checked 25/4 -2025). For bedMethyl files with the preprocessed HG002 data from chromosome 1 and the BSseq object of chromosome 24 from the nine silversides samples, data can be obtained via sharelink from the corresponding author. The code and documentation for the functions is implemented in BSseq (>=1.45.1) available at https://github.com/hansenlab/bsseq/ while we expect a stable version with extended documentation to be included in the next bioconductor release (3.22).

## Acknowledgements

We thank Shyam Gopalakrishnan, Winston Timp, Adam Davidovich, Carolina Montano, Luke Morina, Kaan Okay, Carter Norton, and Yixuan Chen for inspiring discussions on the topic. We also thank Nina O. Therkildsen, Harmony Borchardt-Wier, Maria Akopyan, and Jessica Rick for samples, illustrations and extracting DNA from Atlantic silversides. This project was funded by The Danish National Research Foundation (CEH – DNRF143) and a Carlsberg Foundation award to M.T.L. (CF21-0356).

## Extended Data

**Table 1:** Precision of .99 filtered Nanopore data.

| True CpG status | .99 filtered Nanopore data |  |  |  |
| --- | --- | --- | --- | --- |
|  | Homozygous CpG | Heterozygous CpG | Non-CpG | All CpG |
| Homozygous reference CpG | 1,987,685<br>99.471% | 10,575<br>0.529% | 0<br>0.000% | 2,080,168<br>100.000% |
| Heterozygous reference CpG | 61<br>0.237% | 25,597<br>99.599% | 42<br>0.163% | 25,833<br>99.838% |
| *Non-CpG | 0 | 986 | 810,052 | 1,001 |
| Homozygous non-reference CpG | 16,137<br>98.444% | 249<br>1.519% | 6<br>0.037% | 17,035<br>99.965% |
| Heterozygous non-reference CpG | 12<br>0.055% | 21,685<br>99.733% | 46<br>0.212% | 21,827<br>99.790% |

**Table 2:** Precision of .99 filtered PacBio data.

| True CpG status | .99 filtered PacBio data |  |  |  |
| --- | --- | --- | --- | --- |
|  | Homozygous CpG | Heterozygous CpG | Non-CpG | All CpG |
| Homozygous reference CpG | 1,894,962<br>99.844% | 2,967<br>0.156% | 0<br>0.000% | 2,079,323<br>100.000% |
| Heterozygous reference CpG | 46<br>0.187% | 24,557<br>99.809% | 1<br>0.004% | 25,453<br>99.996% |
| *Non-CpG | 1 | 70 | 13599 | 82 |
| Homozygous non-reference CpG | 15,543<br>99.057% | 148<br>0.943% | 0<br>0.000% | 17,010<br>100.000% |
| Heterozygous non-reference CpG | 16<br>0.076% | 20,942<br>99.900% | 5<br>0.024% | 21,528<br>99.977% |

**Table 3:** Precision of joint .99 filtered Nanopore and PacBio data.

| True CpG status | joint .99 filtered Nanopore and PacBio data |  |  |  |
| --- | --- | --- | --- | --- |
|  | Homozygous CpG | Heterozygous CpG | Non-CpG | All CpG |
| Homozygous reference CpG | 1,818,514<br>99.988% | 216<br>0.012% | 0<br>0.000% | 2,077,928<br>100.000% |
| Heterozygous reference CpG | 21<br>0.087% | 23,981<br>99.913% | 0<br>0.000% | 24,915<br>100.000% |
| *Non-CpG | 0 | 21 | 1,000 | 23 |
| Homozygous non-reference CpG | 14,895<br>99.221% | 117<br>0.779% | 0<br>0.000% | 17,003<br>100.000% |
| Heterozygous non-reference CpG | 6<br>0.029% | 20,419<br>99.971% | 0<br>0.000% | 21,066<br>100.000% |

**Table 4:** Reference and non-reference CpG loci in joint .99 filtered Nanopore and PacBio data.

|  | .99 all filtered |  | .99 homozygous filtered |  | .99 heterozygous filtered |  |
| --- | --- | --- | --- | --- | --- | --- |
|  | reference | non-reference | reference | non-reference | reference | non-reference |
| Total loci | 2,109,124 | 39,017 | 1,823,245 | 15,040 | 24,739 | 21,034 |
| Loci in truth set | 2,102,843 | 38,069 | 1,818,514 | 14,895 | 23,981 | 20,419 |
| Loci not in truth set | 6,281 | 948 | 4,731 | 145 | 758 | 615 |

**Table 5:** Reference and non-reference CpG loci in joint .99 filtered Nanopore data from Atlantic silverside.

|  | .99 all filtered |  | .99 homozygous filtered |  | .99 heterozygous filtered |  |
| --- | --- | --- | --- | --- | --- | --- |
|  | reference | non-reference | reference | non-reference | reference | non-reference |
| RR | 304,363 | 13,658 | 233,947 | 3,202 | 4,281 | 2,410 |
| RN | 299,735 | 28,220 | 206,720 | 3,093 | 18,320 | 16,298 |
| NN | 282,525 | 31,743 | 209,379 | 16,667 | 3,341 | 2,391 |

**Table 6:** Mean methylation of reference and non-reference CpG loci in joint .99 filtered data from Atlantic silverside.

|  | .99 all filtered |  | .99 homozygous filtered |  | .99 heterozygous filtered |  |
| --- | --- | --- | --- | --- | --- | --- |
|  | reference | non-reference | reference | non-reference | reference | non-reference |
| RR1849 | 65.3% | 71.1% | 64.9% | 70.5% | 74.4% | 71.9% |
| RR1871 | 64.9% | 70.9% | 64.5% | 70.1% | 74.1% | 71.8% |
| RR1881 | 65.2% | 70.7% | 64.8% | 70.0% | 73.6% | 71.4% |
| RN1841 | 64.2% | 68.0% | 64.0% | 68.8% | 67.9% | 66.7% |
| RN1860 | 66.5% | 70.2% | 66.2% | 71.0% | 70.0% | 69.1% |
| RN1860 | 65.7% | 69.4% | 65.4% | 70.4% | 69.5% | 68.2% |
| NN1854 | 66.5% | 70.3% | 66.4% | 69.5% | 75.6% | 73.8% |
| NN1867 | 65.8% | 69.3% | 65.7% | 68.7% | 74.7% | 72.9% |
| NN1870 | 64.3% | 67.4% | 64.2% | 66.8% | 72.9% | 70.8% |

**Table 7:** Mean methylation of reference and non-reference CpG loci in .99 all filtered data from HG002.

|  | .99 all filtered |  | .99 homozygous filtered |  | .99 heterozygous filtered |  |
| --- | --- | --- | --- | --- | --- | --- |
|  | reference | non-reference | reference | non-reference | reference | non-reference |
| Nanopore | 67.9% | 70.2% | 69.5% | 71.9% | 71.5% | 69.7% |
| PacBio* | 65.7% | 66.6% | 66.8% | 67.5% | 67.7% | 66.2% |

\* The methylation calls from PacBio are not quality filtered and not comparable with the Nanopore estimates

**Extended data: Fig 1:**
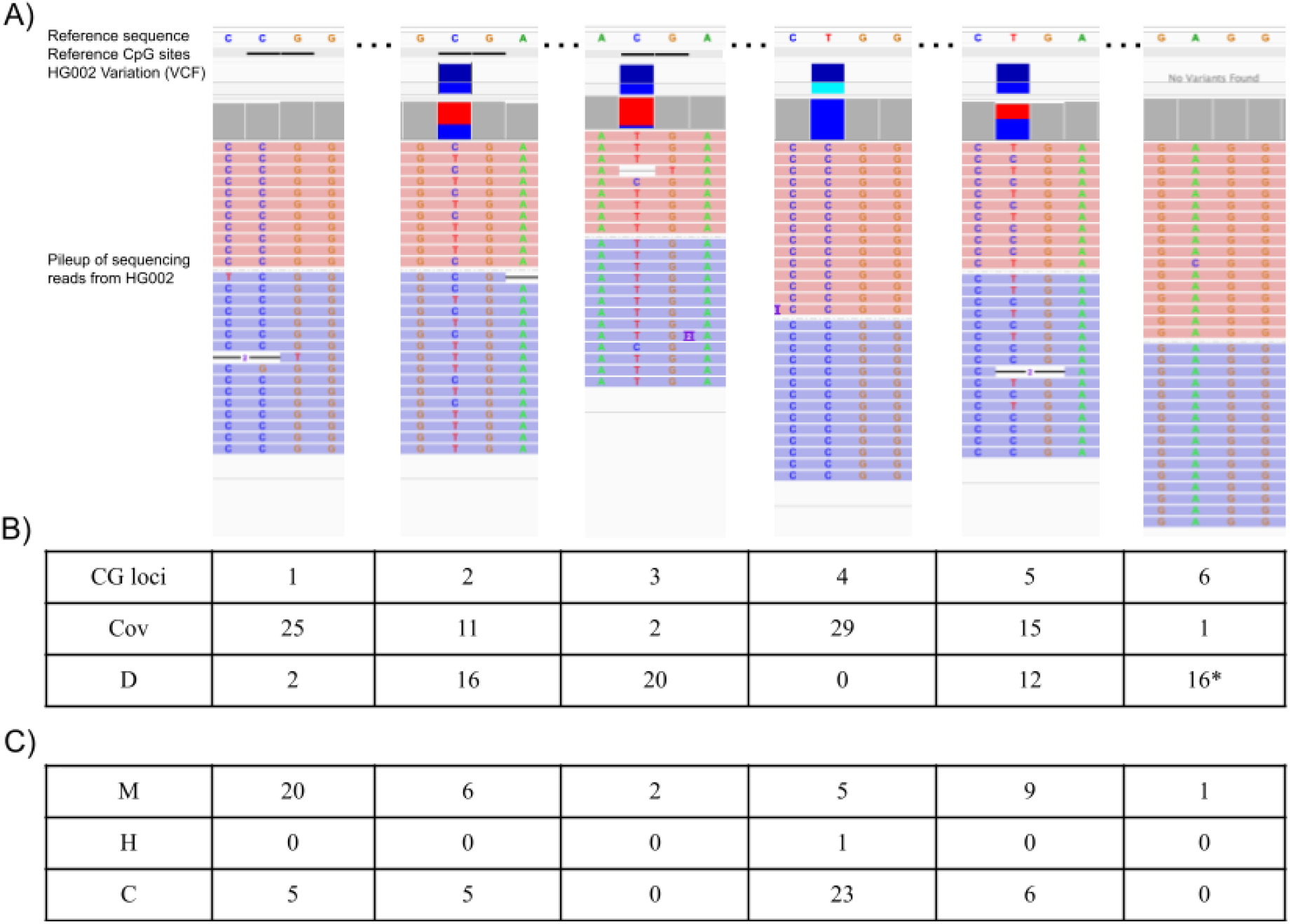
Summarizing nucleotide and modification information from long-read sequencing. **A)** IGV representation of ONT sequencing reads from HG002 mapped against the GRCh38 reference genome assembly. Each row represents a mapped sequencing read, where red colored reads are mapped to the forward strand represented in the reference sequence, and blue colored reads are mapped to the reverse strand. High quality variant calls for the sample are represented in the HG002 Variation (VCF). **B)** The information of CpG and non-CpG coverage per loci from a bedMethyl file imported into BSseq using read.bedMethyl(). Cov represent the number of times a CpG site is mapped to the loci, while D represent the sum of non-CpG sites mapped to the loci ie. deletions (represented by - in A), and different calls than C in CpG context. (*) No Cs in CpG context are detected on the reverse strand, wherefore the information from the reverse strand is not included in the bedMethyl file or in BSseq. **C)** The modification calls from the CpG sites imported into the BSseq object as M, for 5mC, H, for 5hmC and C, for canonical cytosine.

**Extended data: Fig 2:**
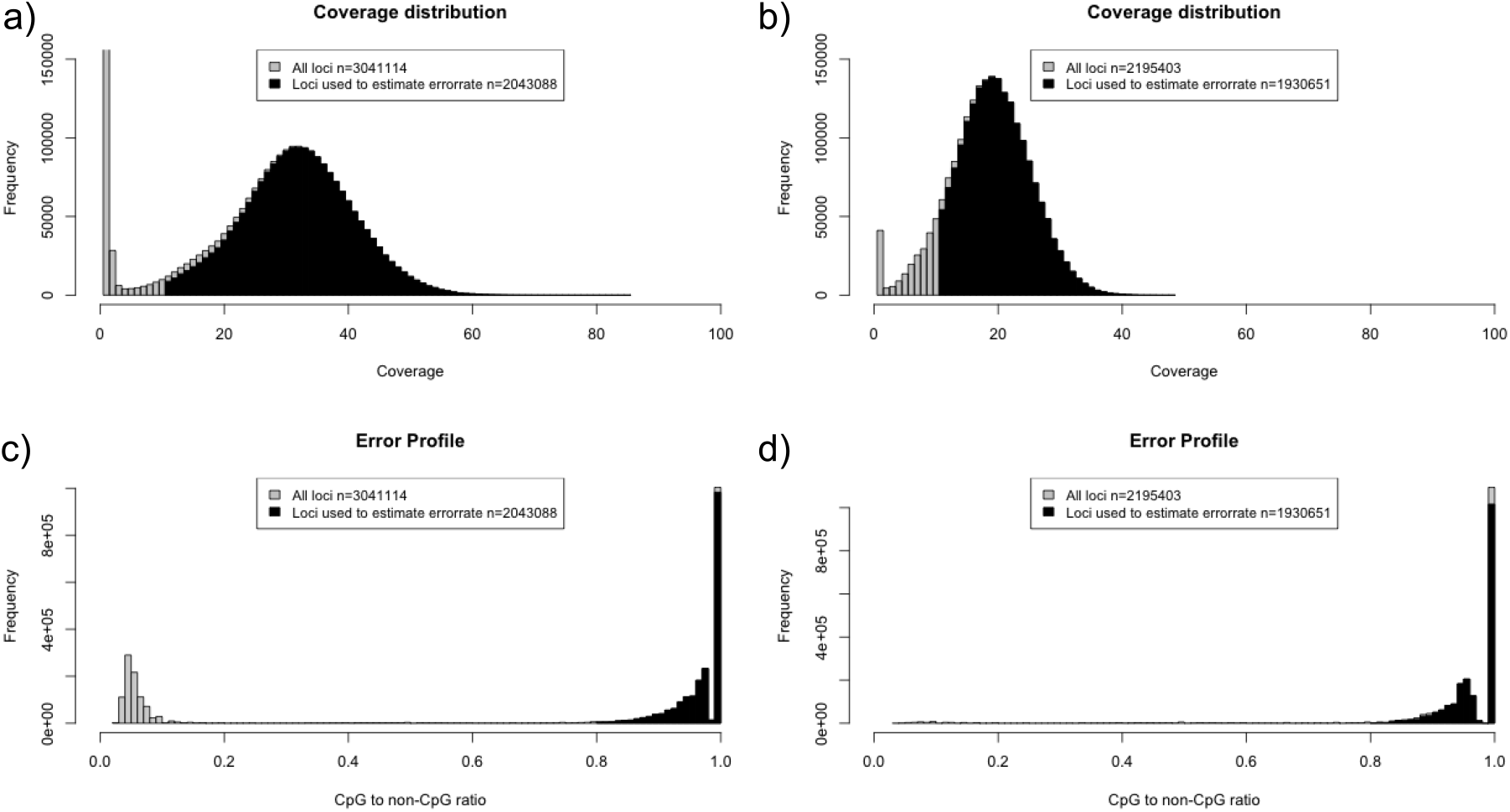
Estimating CpG-specific error rate from long-read sequencing data. **a-b)** The CpG coverage of all loci in the BSseq object in gray and the loci used to estimate the error rate overlaying in black for the Nanopore sample **(a)**, and PacBio sample **(b)** . **c-d)** The CpG coverage divided by the non-CpG coverage of all loci in the BSseq object in gray and for sites used to estimate the error rate in black for the Nanopore sample **(c)** and PacBio sample **(d)**.

## Supplementary methods

### Summarizing nucleotides and modifications from long-read sequencing data

The long-read sequencing data produced by the platforms from Nanopore and PacBio contain information of both the sequenced nucleotides and their modification state, if base called using a modification model. Here, we focus exclusively on 5mC in the CpG context. The standard workflow following base and modification calling is to map the reads against a reference genome, whereafter mapped reads can be examined in pileup format e.g. using IGV^23^(Extended data: Fig. 1a). The reads contain the modification state every time the nucleotide context CpG is sequenced, independent of the reference genome. This information is retained in the bedMethyl files produced by the pileup function in the software Modkit developed and maintained by Nanopore^22^. Consequently, if any modification model for CpG context is used for modification and base calling of Nanopore or Pacbio data, the bedMethyl file produced by Modkit can be used to identify sample-specific CpG loci independent of the reference genome.

To obtain the information for identification of sample-specific CpG loci, we retain the count of reads supporting that a reference locus is respectively a CpG locus (Cov), or a non-CpG locus (D). The count of reads supporting that a locus in the reference genome is a CpG is reported in the N_valid_cov column in the bedMethyl file. And if the CpG count for a reference locus is at least one, the number of reads supporting the locus is a non-CpG (D) is available as sum the of the columns N_delete, N_diff, and N_nocall (Extended data: Fig 1b) ^22^. To remove noise, many workflows use reference-guided filtering by setting the flag --cpg or --traditional-preset to only pile up modification information at positions where the reference locus is CpG. Contrary, no reference-guided filtering or direct input of bam files outputs data for all positions with a CpG coverage of at least one, including noise from sequencing errors^24^.

### Estimating CpG-specific error rates

The error rate in the mapped sequencing reads is an important parameter to distinguish signal from noise. In this context we refer to the error rate as the rate at which true CpG sites are classified as non-CpG sites and vice versa, which can be caused by both sequencing, base calling, and mapping errors. The sequencing and base calling error rate depends on the sequencing technology and is approximated in the phred-scaled base quality string in bam files. Mapping errors, where a read is erroneously mapped to another region than it originates from, depends not only on the quality of sequencing and basecalling, but also the reference genome, and the mapping program. These parameters often vary across experiments and samples and we propose to approximate the mean error rate from the bedMethyl file, using the number of non-CpG sites mapped to homozygous CpG loci. Specifically, we filter for loci with a high probability of true being homozygous CpG and estimate the error rate from the number of non-CpG calls mapping to these using Equation 1.

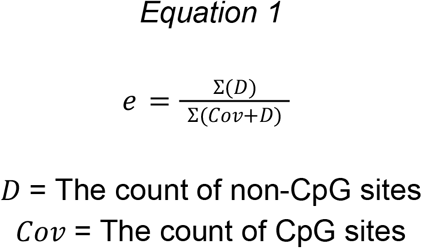

### Computing the scaled likelihood of the data given the CpG status

The likelihood of the data given the error rate and CpG status of a locus can be computed using likelihoods. Here we use a modified version of the genotype likelihood function from GATK^25^ to compute the likelihood of the data (count of CpG and non-CpG) given the loci is a homozygous CpG, a heterozygous CpG or a non-CpG.

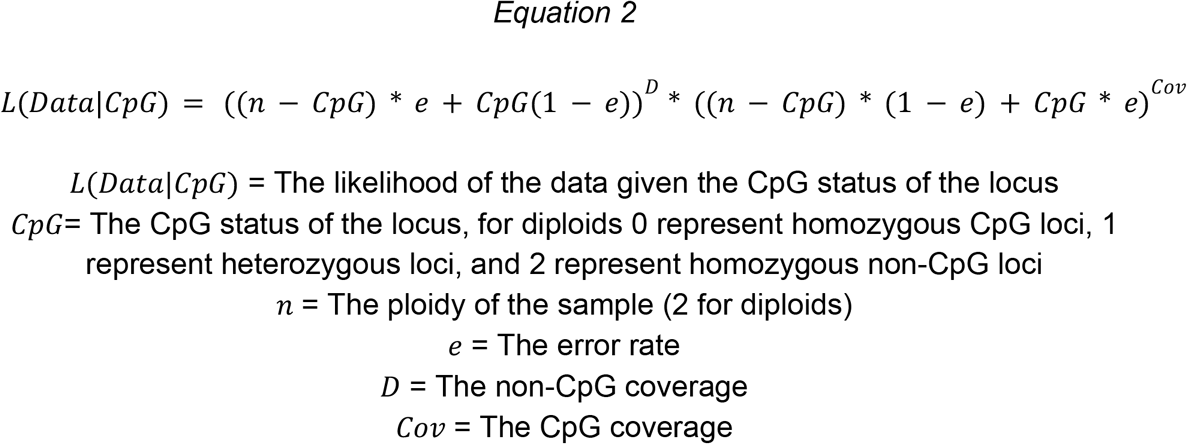

We scale *L*(*Data*|*CpG*) for each locus to sum to one, under the assumption that a locus is either homozygous CpG, heterozygous CpG, or non-CpG.

### Implementation in BSseq

We have implemented functions to read bedMethyl files, compute likelihoods and filter one or multiple samples using the framework presented above in the R-package BSseq. We briefly describe the functions added here, while the full documentation and usage examples are available in BSseq v.1.45.1:

**read.bedMethyl()** reads bedMethyl file(s) generated by Modkit into R as a BSseq object. The BSseq object include CpG counts (Cov) as the sum of canonical C (C), methylated C (5mC, M), and if present hydroxymethylated C (5hmC, H), as well as non-CpG counts (D).

**estimateErrorRate()** uses CpG counts and non-CpG counts from loci with a high probability of being homozygous CpG loci to estimate the error rate using *equation 1*.

**getScaledLikelihoods()** uses *equation 2* to output the scaled likelihood of the data (CpG and non-CpG count) given the CpG status (homozygous CpG, heterozygous CpG or non-CpG) and error rate.

**getCpGs()** outputs the loci with a scaled likelihood above a threshold (default = 0.99) of being either homozygous CpG, heterozygous CpG, homozygous or heterozygous CpG, or non-CpG depending on the type arguement.

**getCpGMatrix()** outputs the most likely CpG status for each loci of each sample in a single or multi-sample BSseq object, which can be used for filtering or genotypic analyses.

**getMaxLikelihoodMatrix()** outputs the scaled likelihood of the most likely CpG status outputted by **getCpGMatrix()**.

### Vocabulary

Base calling: Determining the sequenced nucleotide {A,C,G or T} from the raw sequencing signal.

Modification calling: Determining the modification state of the nucleotide from the raw sequencing signal. Here mainly {C or 5mC} for C in CpG sites.

CpG site: A cytosine followed by a guanine in a DNA read or strand from 5’ to 3’.

CpG locus: The combined term for the CpG site the forward and reverse strand in double stranded DNA.

CpG calling: Determining the CpG status {Homozygous CpG, heterozygous CpG or non-CpG} of a loci in a mapped sample relative to the position in the reference genome.

.99 homozygous CpG filtering: Filtering for loci with scaled likelihood above 99% of being a homozygous CpG locus given the data.

.99 heterozygous CpG filtering: Filtering for loci with scaled likelihood of above 99% of being a heterozygous CpG locus given the data.

.99 all CpG filtering: Filtering for loci with scaled likelihood of above 99% of being a homozygous CpG locus or heterozygous CpG locus given the data.

Joint filtering: Filtering for loci with a scaled likelihood above a threshold for the same CpG status in multiple samples. E.g. joint .99 heterozygous CpG filtering of the Nanopore and PacBio sample gives the loci from .99 filtering in both samples.

Reference-guided filtering: Using only the loci in a sample or sample set which overlap reference CpG loci (setting --cpg in Modkit).

